# Investigation of vitamin B_12_ concentrations and tissue distributions in larval and adult Pacific oysters and related bivalves

**DOI:** 10.1101/2021.10.08.463682

**Authors:** Susanne Vogeler, Gary H. Wikfors, Xiaoxu Li, Justine Sauvage, Alyssa Joyce

**Author notes:** Corresponding author: Alyssa Joyce. **Authors’ contribution** Experimental design and conceptualization were generated by SV, AJ, JS & XL. All laboratory studies and experiments were completed by SV & JS. Manuscript preparation and interpretation of data were conducted by SV, AJ, GHW, and all authors have read and approved the final manuscript.

## Abstract

Vitamin B_12_ (B_12_) is an essential micronutrient for all animals, but is not present in plants and is produced *de novo* only by bacteria or archaea. Accordingly, humans must derive required B_12_ from eating animal products or vitamin supplements, as deficiencies can lead to severe health issues including neuropathy. An often overlooked source in the human diet of B_12_ is shellfish, in particular bivalves, which have significantly higher levels of B_12_ than other animal sources, including all vertebrate meats. Origins and key metabolic processes involving B_12_ in bivalves remain largely unknown, despite the exceptionally high levels. In this study, we examined in several Australian bivalve species, hypotheses concerning B_12_ utilisation and uptake through diet or microorganism symbiosis. Vitamin B_12_ is not distributed evenly across different tissues types of the Pacific oyster, the commercial scallop and Goolwa cockle (pipi), with higher accumulation in the oyster adductor muscle and gill, and mantle and syphons of the Goolwa cockle. Oyster larvae before first feeding already contained high amount of B_12_; however, a significant decrease in B_12_ concentration post metamorphosis indicates a higher utilisation of B_12_ during this life event. We demonstrated that microalgal feed can be supplemented with B_12_, resulting in an enriched feed, but this did not result in an increase in larval B_12_ concentrations when oyster larvae were fed with this diet relative to controls, thus supporting the theory that a B_12_ producing microbiome within bivalves was the potential source of B_12_ rather than feed. However, B_12_ concentrations in the digestive tract of adult oysters were low compared to other tissue types, which might challenge this theory, at least in adults. Our findings provide insight into B_12_ uptake and function in bivalve species, which will aid the promotion of bivalves as suitable B_12_ source for humans as well as provide crucial information to the aquaculture industry in relation to optimisation of vitamin supplementation in bivalve hatchery production.

## Introduction

Vitamin B_12_ (B_12_), or cobalamin, is an essential vitamin for metazoan species that is required in key metabolic processes such as DNA synthesis and fatty acid and amino acid metabolism, as well as playing a functional role in the nervous system (1). Deficiency in B_12_, can cause serious health conditions in humans, including pernicious anaemia, peripheral neuropathy, and other neurological complications (1, 2). Vitamin B_12_ is produced *de novo* in prokaryotes - bacteria and archaea - using aerobic and anaerobic pathways, but no eukaryotes are known to produce this essential vitamin (3-5). In humans, dietary sources of B_12_ include animal products such as meat and dairy (1, 6) (recommended for adults approx. 2.4 µg per day (7)). All carnivores and omnivores must acquire B_12_ from the diet as they have lost cellular synthesis pathways; however, some herbivores have symbiotic relationships with bacteria inhabiting the digestive tract, e.g., ruminants such as cattle, that produce B_12_ *in situ* (8).

A widely underappreciated source for B_12_ for humans is shellfish. Bivalves such as clams and oysters are known to have considerably higher B_12_ concentrations than meat, and therefore provide an excellent natural food-based source of B_12_. Concentrations of B_12_ in bivalves range from 15 µg/100g to 96 µg/100g (9-13), which is much higher than in commonly consumed animal products, such as beef (0.7-5.2 µg/100g), pork (0.4-2.0 µg/100g), and chicken (0.2-0.6 µg/100g) (6).

The reason for such high B_12_ concentrations in shellfish is unknown. In humans and many vertebrates, B_12_ is involved in a wide variety of metabolic functions, including in the circulatory and nervous systems, thus it is surprising that shellfish have such high levels given they have neither red blood cells nor a complex nervous system. Furthermore, it is unclear how bivalves obtain such high B_12_ concentrations as bivalves cannot produce B_12_-only prokaryotes can - thus they must acquire it either through their diet by feeding on microalgae or bacteria that contain it, obtaining the vitamin from microorganisms from their microbiome or by assimilating dissolved B_12_.

Approximately half of the eukaryotic microalgae are dependent on exogenous B_12_ sources (B_12_ auxotrophs) (14); these most often are thought to obtain B_12_ through symbiotic relationships with heterotrophic bacteria (15-19). Algal strains, which are commonly grown for aquaculture diets, such as *Tetraselmis* spp., *Pavlova* spp. and *Isochrysis* spp., contain approximately 180 µg/100g up to approximately 700 µg/100g B_12_ dry weight (20, 21), thus providing an adequate source for B_12_ for bivalves. Bivalves might also assimilate B_12_ from ingested bacteria that are part of their feed, even though algae are considered their primary feed source.

Alternatively, bivalves might also assimilate B_12_ from microorganism in their gut similar to other animals such as terrestrial ruminants (e.g., cattle, sheep) that possess a B_12_ producing microbiome. Such relationships may be particularly important in early larval stages when bivalve larvae first acquire a gut microbiome by ingesting microorganisms from the environment (22).

Although it is possible that bivalves can assimilate dissolved B_12_ directly from seawater, the high quantities in tissue make it seem improbable that this is the primary source. Dissolved B_12_ in seawater is naturally very low, for example, ranging from 0.2-990 pM (23) or 3-605 ng/L in coastal areas of the Northern and Southern Atlantic Ocean (24). Nevertheless, understanding which of these pathways – feed, microbiome, or dissolved vitamins – is responsible for the high concentrations of B_12_ in bivalve species could have practical implications for optimization of rearing conditions to ensure optimal bioavailability of this key nutrient in hatchery production. More globally however, insights into B_12_ levels in different bivalve species may also lead to a better scientific understanding of the role of this vitamin, including its production, uptake and metabolism. A focus on the high levels of B_12_ in bivalves may also help to promote their health benefits when marketing shellfish products.

Vitamin B_12_ deficiency is of concern in human medicine where inadequate intake and malabsorption (9, 25) lead to B_12_ deficiencies, even in countries with adequate access to animal-sourced protein. Studies have shown that B_12_ deficiency occurs in a wide range of age groups, but can be particularly important in children, women of reproductive age, and the elderly (9, 26, 27). The most ubiquitous B_12_ available from natural sources are adenosylcobalamin (AdCbl), methylcobalamin (MeCbl) and hydroxocobalamin (OHCbl) (28). AdCbl and MeCbl are biologically active forms of B_12_, but OHCbl and the industrially-produced cyanocobalamin (CnCbl), which is the form commonly used in nutritional supplements, have to be metabolized to be used by humans (28, 29). Indeed, food fortification and supplementation with MeCbl or synthesized CnCbl can aid against the emerging B_12_ deficiencies; however, recent investigations have challenged the suitability of B_12_ supplements in relation to how well humans are able to uptake specific types of synthetic B_12_ (CnCbl), as well as absorption limitations and genetic conditions that interfere with vitamin B_12_ assimilations (30). For these reasons, natural sources of B_12_ such as occur in shellfish are considered preferable to supplements.

No theory has been advanced as to why bivalves have such high levels of B_12_. Molluscs including most bivalve species have significantly higher levels of B_12_ than other phyla of marine invertebrates (31) and much higher B_12_ levels than fish or terrestrial livestock animals. In an attempt to answer why, in this study, we explored B_12_ distribution in several adult bivalve species and investigated B_12_ assimilation during different developmental stages of oysters. We experimentally varied B_12_ during oyster larval development and assessed whether or not high B_12_ diets (delivered as B_12_-enriched microalgae) affected the B_12_ concentrations in larvae. We also examine the B_12_ concentrations in species of microalgae commonly used in the aquaculture industry when grown in differently B_12_-enriched growth media in order to assess the effects of different B_12_ levels in feeds. Information derived from these experiments provides a preliminary overview of the utilisation and distribution of this key vitamin.

## Materials and Methods

Algal cultivation and larval experiments were carried out at the South Australian Research and Development Institute (SARDI) in Adelaide, Australia.

### Vitamin B_12_ in bivalve tissues

Adult Pacific oysters, *Crassostrea gigas*, were obtained from Franklin Harbour, South Australia, including the parental generation for the spawned larvae that were utilised in subsequent larval experiments. From a local fishmonger, we purchased alive Goolwa cockles, *Plebidonax deltoides* (‘pipis’) also from South Australia, and commercial scallops *Pecten fumatus* (adductor muscle and roe) sourced from Victoria, Australia. The oyster samples were sexed prior to dissection, and the following tissue types were sampled from three males and three females: mantle (one side), gills (one side), posterior adductor muscle, gonad and digestive tract (digestive gland, stomach, midgut, intestine). The adductor muscles from commercial scallops, with roe attached, were dissected after defrosting. The tissue of a single (logistical limitations) female Goolwa cockle was also dissected: foot, adductor muscle, gonads (eggs), digestive tract and remaining parts (gills, mantle and siphons). Each tissue sample was rinsed in fresh tap water to remove debris, homogenised using a hand blender, and stored at -80°C for B_12_ analysis. Weight of sampled tissues ranged from 0.399 g to 1.434 g wet weight, depending upon tissue type (S1 File)

### Algal feed cultivation

Four species of microalgae, commonly used in bivalve hatcheries and known to be B_12_-dependent, were used in the feeding experiments: *Tisochrysis lutea (T-iso), Pavlova lutheri, Chaetoceros muelleri*, and *Chaetoceros calcitrans* (19, 32). All species were grown separately in UV-treated, 0.04-µm-filtered seawater (38 ppt, pH 8.2) and provided with 24 h light and aeration with additional CO_2_. Algae were cultivated in different types of media (S2 File): (1) f/2 medium (33) with a final low B_12_ concentration of 3.69×10^−10^ M (1xB_12_), (2) f/2 with a final high B_12_ concentration of 3.69×10^−9^ M (10xB_12_), or (3) Walne medium (34) with a final B_12_ concentration of 7.38×10^−9^ M (20xB_12_). Microalgae grown in f/2 media were cultured under static conditions in 10-L carboys. Stock solutions for essential vitamins B_12_, thiamine (B_1_), and biotin (formerly vitamin H, now B_7_) were prepared in 0.2-µm-filtered seawater and added to the f/2 media after autoclaving. Microalgae grown in Walne medium were cultivated by the SARDI hatchery staff with *T. lutea, P. lutheri* and *C. muelleri* maintained separately in 50-L, semi-continuous culture bag systems. The final Walne medium for the bag system was heated to 80°C for pasteurization and cooled before being added to the algal cultures. *C. calcitrans* cultures grown in Walne medium were cultivated in 10-L carboys with the final media autoclaved before use.

Algal species grown in 1xB_12_ f/2 medium or 10xB_12_ f/2 medium, as well as in Walne medium (20xB_12_) were sampled for B_12_ analysis from three different carboys or bags per species/medium. One additional sample for algae grown in the Walne medium (continuous bag culture) was taken each for *T. lutea* and *P. lutheri*. Approximately 1 L to 1.5 L per carboy or bag of each algal culture was harvested and centrifuged at 1,960 g for 2 min to pellet the algae. Carboys were sampled at a density of 5.1×10^6^ – 1.5×10^7^ cells/ml; whereas, bags at exponential growth phase were sampled by hatchery staff. After centrifugation, 50 mL of the original supernatant for each algal sample was collected for analysis. Algal pellets were resuspended in 50 mL fresh seawater as an additional washing step, then centrifuged again (the second supernatant was discarded). The algae pellets and media samples were stored at -80°C. For the B_12_ analysis, all algal samples were freeze dried and stored at -80°C. Dried larval pellets weight ranged from 0.041 g to 0.198 g (S1 File).

Additional media samples were collected from fresh 1xB_12_ f/2 medium, 10xB_12_ f/2 medium, autoclaved Walne medium, non-autoclaved Walne medium, and pasteurized (80°C) Walne medium from the bag system. For each media type, three 50 mL samples were taken and frozen at -80°C for further analysis.

### Feeding experiments

Pacific oyster, *C. gigas*, larvae were derived from fourteen family lines of broodstock originating from Franklin Harbour in South Australia. Larvae were fed the first time 24 hours post fertilisation (hpf), when larvae had reached D-shelled larval stage, with daily feedings thereafter with microalgae grown either in 1xB_12_ f/2 medium, 10xB_12_ f/2 medium or Walne medium. The first 7 days post fertilisation (dpf), oyster larvae were fed with a microalgal mix consisting of *T. lutea, P. lutheri* and *C. calcitrans*. Both, *T. lutea* and *P. lutheri*, were given at equal ratios and the feed density was increased gradually each day from 30,000 cells/mL to 50,000 cells/mL, while *C. calcitrans* was fed at a constant volume corresponding to 20,000 cells/ml. From the eighth day onwards, *C. muelleri* was added to the feed mixture when larvae reached a shell size at which this species is known to be ingested. The mixture then consisted of 30% *T. lutea*, 30% *P. lutheri*, and 40% *C. muelleri*. The feed density was increased each day until 14 dpf to 80,000 cells/mL and kept constant until end of the experiments, while *C. calcitrans* was given at a constant volume corresponding to 20,000 cells/ml.

### Low and high vitamin B_12_ diets during larvae development

Prior the first feeding, 24 hpf D-shelled larvae were placed in 20-L conical tanks and reared under static conditions with gently-aerated, UV-treated, 1-µm-filtered seawater (38 ppt, pH 8.2). The starting stocking density was 8-10 larvae/mL, which was gradually reduced to <1 larva/mL at the end of the experiment as larvae were sampled and graded. The experiment was carried out for 21 dpf until larvae reached the late-veliger stage. All larvae were fed daily with an algal mixture grown either in 1xB_12_ f/2 or 10xB_12_ f/2 after each tank was cleaned and refilled with new seawater. Four individual tanks as biological replicates were maintained per algae treatment.

Over the course of the experiment, larval density and size were monitored every 2-4 days and assessed under an inverted microscope. After the experiment was terminated, larvae were washed and kept in seawater without feed for 6 h (depuration). Larvae were then sampled (10,000-20,000 larvae; ∼88-170 mg from all tanks, except for one 10xB_12_ f/2 tank with ∼5,000 larvae (0.016 g)) that were retained through settlement and sampled as spat; in all cases, final size and density were assessed. All larval samples were centrifuged at 1960 g for 3 min to remove remaining seawater and kept at -80°C until further analysis. Three samples were taken at the beginning of the experiment of unfed, 24-hpe, D-shelled larvae, centrifuged as the other samples, and stored at -80°C. Before larval samples were sent for analysis, each sample was homogenised using a hand blender and re-frozen. Temperature of one 20-L tank was assessed continuously over 3 days from 15 dpf to 18 dpf using a temperature sensor.

### Vitamin B_12_ assessment during larval development

To assess the B_12_ concentration throughout development, a second larval experiment was conducted simultaneously using larvae from the same fertilisation event as the feeding experiment. Larvae were reared under the same conditions, but to obtain the larger required volumes for sub-sampling at different developmental stages, larvae in this treatment were instead reared in one 200-L tank under static conditions instead of as replicates of specific treatments in the 20-L tanks. Larvae were fed daily with an algae mixture grown in Walne medium (20x B_12_) as outlined above. Larvae were sampled at different developmental stages, including late D-shelled larvae (3 dpf), early veliger (5 dpf), mid-veliger (10 dpf), late veliger/early pediveliger (15 dpf, before eye-spot developed), and at eyed pediveliger larvae (19 dpf). Larvae displayed the typical behaviour of competence for metamorphosis, including prominent eye-spot, sinking to the bottom, reduced velum, and crawling behaviour (extending the foot) after 19 dpf. Metamorphosis was chemically induced with epinephrine hydrochloride (Sigma-Aldrich) at 10^−4^ M for 1 h (35), and once set, were washed and kept in seawater with algal feed. Spat (21 dpf) were collected and sampled two days post metamorphosis. Both larvae and spat were sampled in triplicates representing three technical replicates and were prepared and stored as described above. Weights of all larval samples were recorded (S1 File). After completion of the experiment, the 200-L tank was filled with seawater and temperature was measured continuously over two days using a temperature sensor.

### Analysis of vitamin B_12_

The B_12_ analysis was conducted by Eurofins Vitamin Testing Denmark, which is an accredited food lab for B_12_ testing using the *Lactobacillus leichmanii* (ATCC 7830) microbiological assay (Reference method: AOAC 952.20). Using this method, B_12_ is extracted from the samples in an autoclave using a buffered solution and then diluted in basal medium. The growth response of *L. leichmanii* to extracted B_12_ is measured turbidimetrically, which is then compared to calibration solutions with known CnCbl. The limit of detection for this assay is indicated as 0.01 µg/100g.

### Statistical analysis

Total B_12_ concentrations for whole oysters were calculated based upon the B_12_ concentrations of each tissue type corrected for the proportion of the total weight. Whole oysters were calculated based upon sampled tissue types (excluding labial palps, heart and connecting tissue, as these were not sampled). Statistical difference between B_12_ concentrations of algae, larvae, and tissue samples were calculated using a Student’s T test for equal or unequal variance detected by an F-Test for Variance or a Kustal-Walis H test followed by pairwise comparison using Dunnett’s T3 method to trimmed means (36).

## Results

### Vitamin B_12_ in adult tissues

Vitamin B_12_ concentrations in five different tissue types -- mantle, gills, digestive tract, adductor muscle, and gonads (eggs/sperm) -- of six *C. gigas* individuals, separated by sex (three males and three females), were assessed (Fig. 1). The B_12_ concentrations of the gill sample of male 1 and the digestive tract and sperm samples of male 3 could not be quantified by the *L. leichmanii* assay (ATCC 7830) conducted by Eurofins and are therefore missing (Fig. 1A). Overall, no significant differences were found between tissue types from male and female individuals; thus mantle, gills, digestive tract and adductor muscle samples of males and females were pooled for statistical analysis (Fig. 1B).

**Fig. 1:**
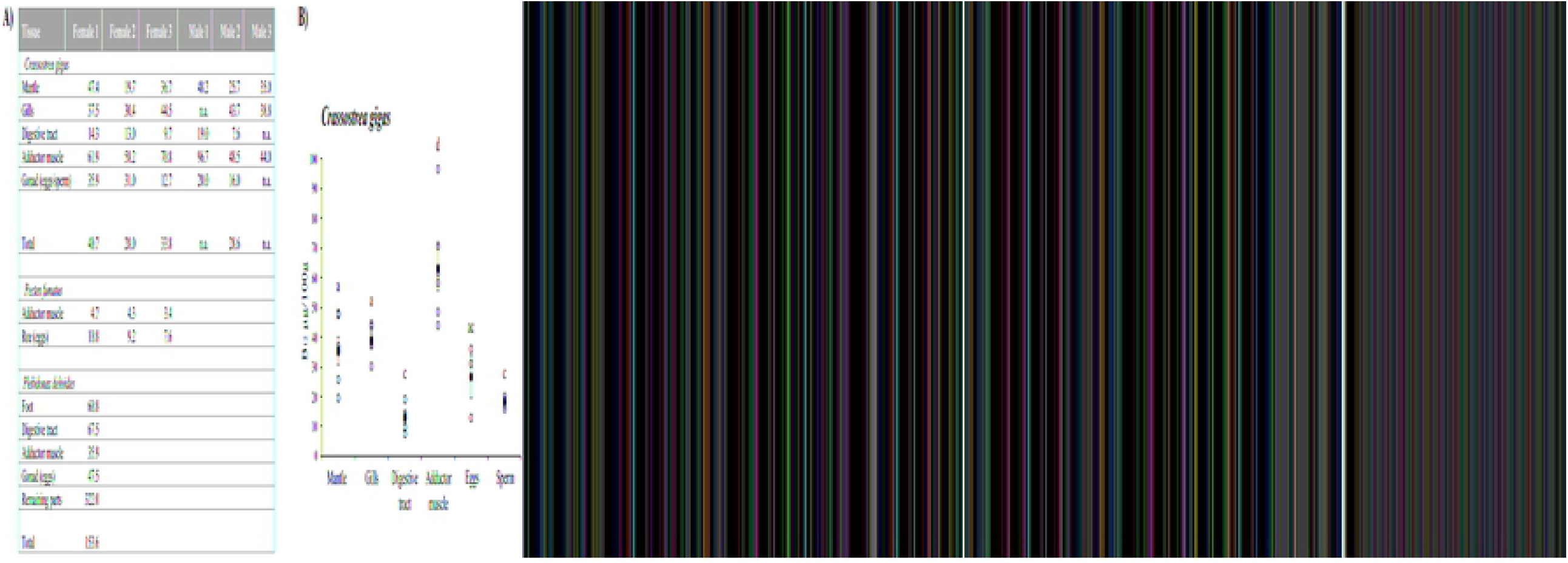
Vitamin B_12_ (µg/100g) concentrations in different tissue types of bivalves. **A**) B_12_ concentration in three female and three male Pacific oyster (*Crassostrea gigas*) individuals, three female commercial scallop (*Pecten fumatus*) individuals and a single female Goolwa cockle (‘pipis’; *Plebidonax deltoides*). n.a: not analysable. Total: total B_12_ concentration corrected for wet weight of tissue to total weight. **B**) Average B_12_ concentration (filled circles) of pooled male and female oyster individuals (open circles) per tissues type. Error bars for average concentration: standard error; letters above show significant differences (p<0.05).

On average, adductor muscle samples contained the highest amount of B_12_ with 63.4±7.7 µg/100g, a significantly higher concentration compared to mantle tissue with 35.5±4.7 µg/100g, gills with 39.0 ±2.5 µg/100g, and female gonads (unfertilised eggs) with 26.5±7.1 µg/100g. The lowest B_12_ concentrations were recorded in male gonads (sperm) with 18.0±2.0 µg/100g and the digestive tract with 12.7±2.0 µg/100g. The total B_12_ concentration for each individual, as a mean of the concentrations in each tissue type in relation to their proportion of total weight, ranged from 28.0 µg/100g to 40.7 µg/100g for the three females and male 2.

In contrast to *C. gigas*, the commercial scallop *P. fumatus* contained significantly lower B_12_ levels (p<0.01) in the adductor muscle with 4.1±0.4 µg/100g. The scallop individuals were bought frozen, and although we cannot exclude that the freezing and storage process might have modified the final B_12_ concentration, single freezing cycles are not noted in literature to affect the final B_12_ concentrations in seawater, blood serum, or fish (37-39). The mean B_12_ concentration in scallop roe, 9.2±0.9 µg/100g, was not significantly different (p>0.05) from unfertilised oyster eggs.

A single female Goolwa cockle, *P. deloides*, individual was also dissected and analysed, which provided a very different distribution of B_12_ in the tissues, with the highest concentrations measured in the remaining parts including the gills, mantle and the siphons (322.0±96.6 µg/100g) followed by the digestive tract with 67.5±20.3 µg/100g, foot with 60.8±18.2 µg/100g, and gonads/eggs with 47.5±14.3 µg/100g. The lowest concentration was in the adductor muscle at 35.9±10.7 µg/100g.

### Vitamin B_12_ in algae

The four algal species all contained B_12_ when grown in 1xB_12_ f/2 (Fig. 2A): *C. muelleri* 98.0 ±1.0 µg/100g > *T. lutea* 71.0 ±2.3 µg/100g > *C. calcitrans* 49.1 ±1.6 µg/100g > *P. lutheri* 38.7 ±1.3 µg/100. Increasing B_12_ concentrations in the media led to significant increases in B_12_ concentration in all algal cultures, with concentrations for algae grown in 10xB_12_ f/2 medium as follows: *T. lutea* 811.0 ±29.8 µg/100 > *P. lutheri* 249.7 ±6.9 µg/100g > *C. muelleri* 202.0 ±4.4 µg/100g > *C. calcitrans* 184.3 ±6.4 µg/100g. When compared to algae grown in the continuous bag cultures in Walne medium, only *P. lutheri* showed a significant increase of B_12_ concentrations with 604.0±136.4 µg/100g; the *T. lutea* bag cultures with 756.5 ±190.2 µg/100g and *C. muelleri* with 301.0 ±49.0 µg/100g were not significantly different from carboys with 10xB_12_ f/2. In contrast to the carboy-cultured algae, however, the algae grown in the bag system displayed large variance between individual samples likely resulting from variation in the growth phase of each culture (concentration of algae in the bag systems varied). As algae in the bag bioreactors were continuous, thus in exponential phase at all times, the B_12_ concentration in each sample varied based upon the concentration of algae in the bag at the time of sampling. Analysis of the supernatant in the algal cultures, however, did not show high variance between samples for *T. lutea, P. lutheri* and *C. muelleri* (Fig. 2B). It needs to be noted that two of the *T. lutea* and *P. lutheri* media samples could not be analysed using the microbiological assay as a consequence of a handling error (Eurofins). In general, media samples for each algal sample displayed a similar pattern in B_12_ concentrations with lower concentrations for 1xB_12_ f/2 medium and significantly higher concentrations in 10xB_12_ f/2 medium, except for *C. muelleri*. Interestingly, the media in the three bag-grown algal species -- *T. lutea, P. lutheri* and *C. muelleri* -- display significantly higher B_12_ concentration than 10xB_12_ f/2 samples, suggesting that the continuous inflow of media in the hatchery bag system led to accumulation of B_12_. Alternatively, it is possible that a general higher starting concentration was delivered in the media supplying the bags as a result of a preference by the hatchery staff for high levels of vitamins and trace minerals in the stock solutions (to counteract unknown effects of pasteurisation). This was confirmed by analysing Walne medium in unused bags (Fig. 2C), which showed significantly higher B_12_ concentrations compared to the unautoclaved Walne medium that was prepared following a standard protocol (S2 File).

**Fig. 2:**
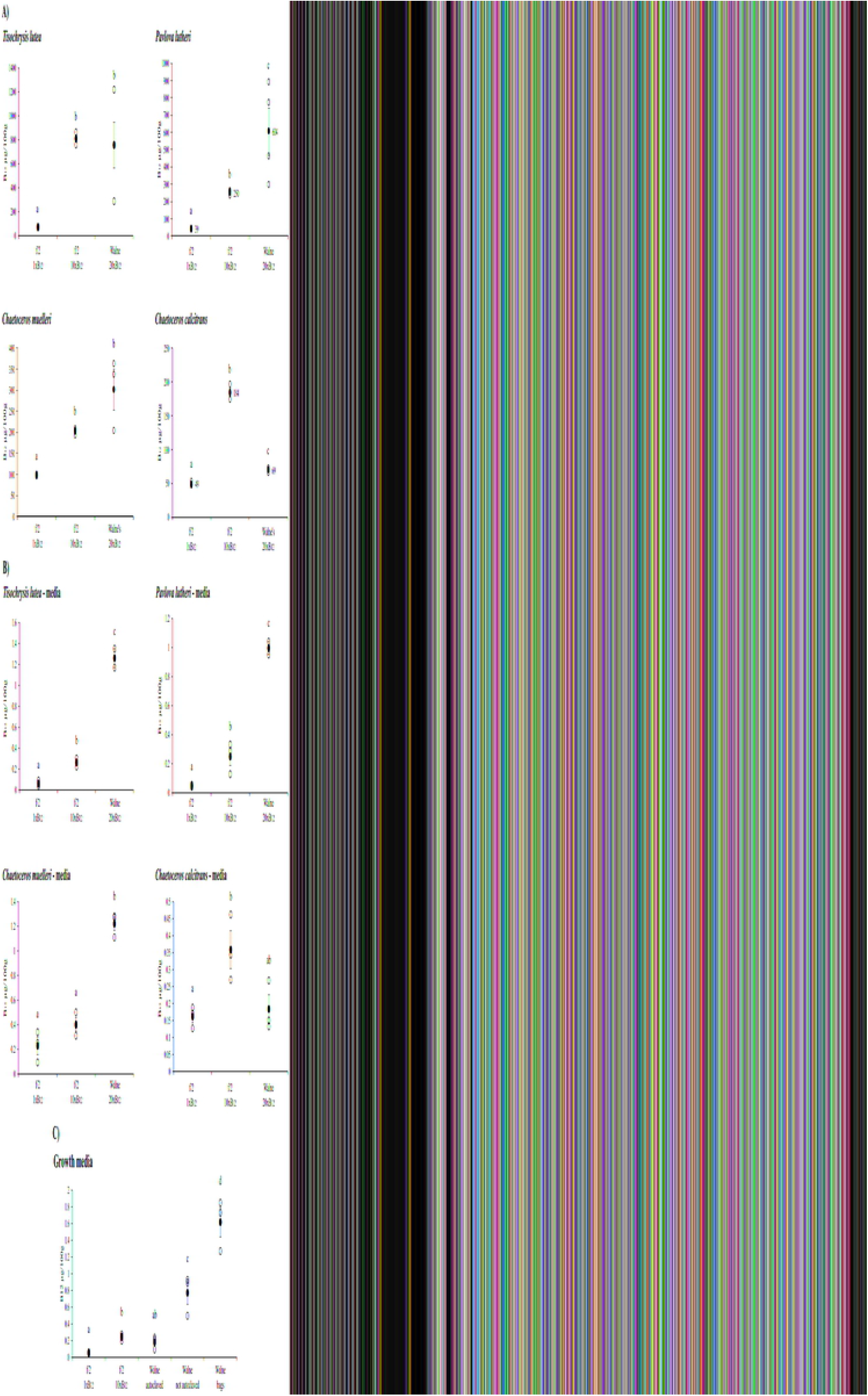
Vitamin B_12_ (µg/100g) concentrations of microalgal species. **A)** Average B_12_ concentrationins in the four microalgal species (*Tisochrysis lutea, Pavlova lutheri, Chaetoceros muelleri* and *Chaetoceros calcitrans;* freeze-dried) grown in 1xB_12_ f/2, 10xB_12_ f/2 and Walne media (20xB_12_) and **B)** of the media supernatant of the four algal species. Algae grown in 1xB_12_ f/2 and 10xB_12_ f/2 media as well as *C. calcitrans* Walne medium were cultivated in 10-L carboys, while the remaining algal species grown in Walne medium were cultivated in a bag system. **C)** B_12_ concentration of fresh growth media for each medium type. Open circles: individual measurements, filled circles: average of all measurement for this sample point including standard error (error bars), letters above show significant differences (p<0.05).

Compared to the other three algal species, *C. calcitrans*, which was cultured exclusively in carboys (batch cultures) rather than the continuous culture bags, displayed a divergent pattern of B_12_ distribution. Although the B_12_ concentrations in algae from 10xB_12_ f/2 were significantly higher than from 1xB_12_ f/2, the cultures grown in Walne medium were significantly lower with 69.2 ±1.6 µg/100g (Fig. 2A). This was also observed in the media supernatant for *C. calcitrans* cultures (Fig. 2B). The carboys with Walne enrichment were autoclaved before algae were added, a practice that is often followed in hatcheries. As we believed the autoclaving to be damaging to the vitamins, we also assessed autoclaved and non-autoclaved Walne media (Fig. 2C). Our results confirm that autoclaving media does indeed significantly reduce B_12_ concentrations, and that autoclaving as standard hatchery protocol to prepare sterile media can significantly destroy vitamins such as B_12_.

### Vitamin B_12_ in oyster larvae

Vitamin B_12_ concentrations were assessed during oyster larval development and after feeding with algae grown in media with different B_12_ concentrations. Larvae at 24 hpf, which were unfed and used for larval feeding experiments, contained a mean B_12_ concentration of 50.0 ±3.0 µg/100g (Fig. 3A). Throughout development, to the end of larval phase (eyed pediveligers), the B_12_ concentrations did not significantly change for larvae fed with algae grown in Walne medium. After metamorphosis, however, mean B_12_ concentration decreased significantly in two-day old spat (21 dpf) to 28.1 ±2.3 µg/100g. The temperature in the 200-L tank varied from 24.40°C to 26.15°C, with an average of 25.38±0.03°C. Larvae reached eyed pediveliger stage and were close to metamorphosis after 19 dpf, with an average size of 317.0±2.3 µm (Fig. 3B).

**Fig. 3:**
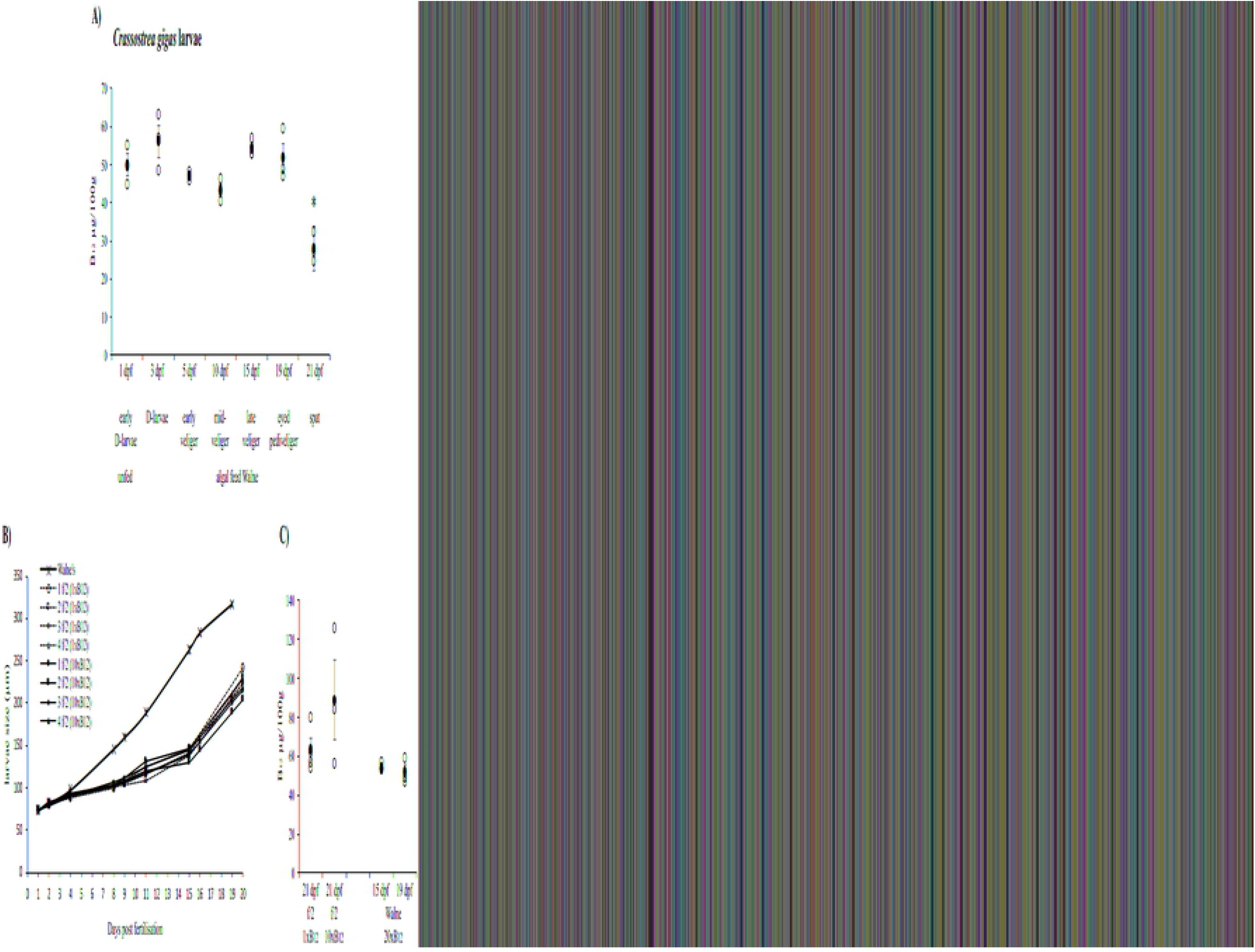
Vitamin B_12_ (µg/100g) concentrations of Pacific oyster, *Crassostrea gigas*, larvae. **A)** Average B_12_ concentrations at different larval live stages. Early D-shelled larvae 1 day post fertilisation (dpf) were assessed prior first feeding and thereafter fed with algal mixtures of algae grown in Walne medium. **B)** Average larval size (µm) during development from 1 day post fertilisation (dpf) until end of experiments with final size assessment on 19 dpf for larvae in one 200 L tank fed with algae grown in Walne medium and on 20 dpf for larvae in each four separate 20-L tanks fed with algae grown in f/2 media (1xB_12_ & 10xB_12_). **C)** B_12_ concentrations of 21 dpf larvae fed with algae mix grown in f/2 medium and 15 dpf and 19 dpf larvae fed with algae mixture grown in Walne medium. Circles: individual measurements, filled circles: average of all measurement for this sample point including standard error (error bars), *: significant different (p<0.05).

Larvae reared in smaller tanks of 20-L volume fed with algae grown in f/2 (1xB_12_ & 10xB_12_) grew significantly slower and did not reach eyed pediveliger stage (Fig. 3B). As there was no significant difference in larval growth between the two B_12_ algal treatments, experiments were terminated when larvae were at late veliger stage (20 dpf, but no predominant foot visible) and reached an average size of 229.8±2.3 µm for larvae fed with 1xB_12_ f/2 algae, and 217.6±2.8 µm for larvae fed with 10xB_12_ f/2 algae. The temperatures in the smaller tanks were lower than the 200-L tank, with an average of 23.30±0.03°C ranging from 22.6°C at night to 25.9°C during the day.

Different B_12_ concentrations in algal feed did not significantly change the B_12_ concentration of oyster larvae (Fig. 3C). Larvae fed with algae grown in f/2 with different B_12_ concentrations did not significantly vary in their final average concentrations after 21 dpf with 1xB_12_ f/2 larvae containing 63.2±5.8 µg/100g and 10xB_12_ f/2 larvae of 88.8±20.30 µg/100g, although the observed trend indicates higher B_12_ levels in the larvae fed with high B_12_ algal feeds. When compared to late veliger (54.4±1.33 µg/100g) and eyed pediveliger larvae (51.8±3.83 µg/100g) fed with algae grown in Walne medium, which contained the highest B_12_ concentration of all algal feed, B_12_ concentrations of 1xB_12_ f/2 larvae and 10xB_12_ f/2 larvae were not significantly different, suggesting that B_12_ concentrations in algal feed does not affect the final B_12_ concentration of larvae at late larval stage, or that the lowest algal B_12_ ration was sufficient or in excess to provide for larval needs.

## Discussion

Shellfish, such as bivalves, are an excellent natural source of bioavailable B_12_ (10, 12, 13, 40, 41). Bivalves have higher B_12_ concentrations than standard meat and dairy products (6) and greater bioavailable B_12_ than other marine species analysed, such as abalone and herbivorous snails (11). Most prior studies have focused on vitamin concentrations in whole animals, with almost no information provided on how B_12_ is distributed in the different tissue types in bivalve species.

The results presented show that in relation to the proportion of the total weight, Pacific oysters tested showed a comparable B_12_ concentration (28-40.7 µg/100g) to whole oysters previously reported (15.1-46.3 µg/100g) (10, 12, 13). The results also indicate that most B_12_ in Pacific oysters is stored in the adductor muscle, accounting for approximately 31-35% of total soft-tissue B_12_; whereas, the lowest concentration of B_12_ was found in the digestive tract. This finding was surprising given that the digestive tract could be presumed to be the organ for intake and absorption of B_12_. Most vertebrates store B_12_ in the liver; in invertebrates, the digestive gland/hepatopancreas is the analogue to the vertebrate liver. In shrimp, for instance, B_12_ is assumed to be stored in the hepatopancreas (42). Thus, it would have been expected to find high levels of B_12_ in the digestive organs of oysters, given that B_12_ could have been contained either in the feed or produced by microbial sources in the gut. These findings are further confounded by the fact that oysters seem to be unique in their storage of B_12_ in the adductor muscle, as the two other bivalve species analysed – scallops and Goolwa cockle – did not exhibit a similar pattern. Vitamin B_12_ concentration in the scallop adductor muscle was significantly lower than that in oysters, with an average of 4.1±0.4 µg/100g. Previous work has reported a B_12_ concentration of 13.4±1.0 µg/100g for the whole body of the scallop species *Mizuhopecten yessoensis* (10) and substantially lower concentrations in adductor muscle at 1.1±0.2 µg/100g (11). Similar methods for B_12_ detection were utilised in those studies, suggesting that the adductor muscle is not the key tissue for B_12_ storage in scallops. The Goolwa cockle also does not appear to store the majority of its B_12_ in the adductor muscle; the adductor muscle had the lowest B_12_ concentrations amongst all tissues tested in this species. Although the digestive tract contained the second highest B_12_ concentration, the majority (77% of whole-body weight) of B_12_ was stored in the remaining cockle soft tissues, such as siphons, gills and mantle tissue. Further research is needed to specify which of the remaining organs contribute to the overall very high B_12_ concentrations observed in the cockle sampled, and reported elsewhere for a wide range of clam/cockle species. Other molluscan species, such as gastropods and edible marine snails, display much higher concentrations in their visceral tissue than in in the edible muscle tissue (11, 43).

The function of such high B_12_ concentrations in bivalves and other molluscan species is speculative. In terrestrial animals, as well as fish, B_12_ is required as a cofactor for two B_12_-dependent enzymes, methylmalonyl-CoA mutase (MCM) for conversion of methylmalonyl-CoA to succinyl-CoA in the mitochondria and methionine synthase (MetH), which catalyses the re-methylation of homocysteine to methionine, an essential amino acid (for review see (44)). Homologues of both B_12_ -dependent enzymes are predicted in the genomes of the Pacific oyster *C. gigas* (protein GenBank ID: MCM (XP_034309642) & MetH (XP_034325685)) and the scallop *P. maximus* (protein GenBank ID: MCM (XP_033762147) & MetH (XP_033734273)), but it is unknown if B_12_ in molluscs functions similarly to key functions in vertebrates, where it is known to be involved in the health of nervous tissue, particularly myelin synthesis (45) and erythropoiesis (46). In marine bivalves, and invertebrates generally, B_12_ might fulfil other functions potentially involved in the immune system. Non-specific immune response, including haemocyte counts, improved at optimum B_12_ concentrations in the juvenile Chinese mitten crab, *Eriocheir sinensis* (47). Many *Vibrio* spp. are thought to be B_12_ scavengers (48); thus a mechanism to remove B_12_ from the gut or gills may also help to control bacterial populations that are scavenging free B_12_. Vitamin B_12_ is known to have a critical role as an antioxidant, and thus may also aid osmotic regulation, which is particularly important for an intertidal species that faces strong salinity fluctuations and long periods during which the shell must remain closed during tidal fluctuations. The high levels of B_12_ in bivalve shellfish could therefore be hypothesised to play a role in oxidative stress responses by reducing homocysteine levels (49, 50).

Although plants do not generally require B_12,_ as they contain an alternative B_12_-independent form of methionine synthase (MetE) (51), bioavailability of B_12_ potentially leads to an increase in phenolic compounds that are able to protect plants against oxidative stress induced by salinity (52). Vitamin B_12_ deficiency also leads to oxidative stress and memory impairment in annelids (53). High demand for methionine production by MetH might also be a unique trait of shell producing animals such as bivalves, given that some bivalves such as pearl oysters contain unique proteins for biocalcification and shell production which are remarkably rich in methionine (54-56). Besides the unknown functions of B_12_ in bivalves, how this crucial vitamin is derived – whether from algal feed or microbiome - has still not been confirmed.

Microalgal species such as *T. lutea, P. lutheri, C. muelleri* and *C. calcitrans*, commonly used in hatcheries as larval feed, are B_12_-dependent (19, 32) and thus high density algal cultures are always supplemented with B_12_ in algal growth media. Haptophytes such as *T. lutea* contain only a MetH homologue (57), but *Chaetoceros* spp. also contain MetH, with some species additionally expressing MetE (58). *P. lutheri*, in addition to MetH, may be able to remodel pseudo-vitamin B_12_ (59), a non-bioavailable form of B_12_ for most eukaryotes that is produced by specific bacteria. In the absence of bacteria which produce B_12_ - such a occurs in axenic algal cultures - microalgae can assimilate B_12_ from the growth media (60, 61). Our results show that B_12_ concentrations in these four microalgal species were significantly increased by providing elevated B_12_ in the growth media. Algae grown in the bag system, however, showed a larger variability in B_12_ concentrations compared to carboy-grown algal cultures, which we cannot fully explain, given we did not see similar variability in the corresponding media supernatants. The bag systems were continuously provided with enrichments, but were also being harvested continually as feed for hatchery production. Consequently, not all sampled bags would have been at similar densities, and drip rates of media and water may have varied slightly, thus influencing the final B_12_ concentrations based on different levels of media and density of cultures. Whether or not there is differential B_12_ uptake during different growth phases is not known, although B_12_ has been shown to be variable in batch cultures in which uptake is highest in the exponential growth phase (57). Levels of B_12_ observed in the diatom *C. calcitrans*, however, which was cultured only in carboys (it does not grow well in hatchery bag systems), demonstrated an important principle related to vitamins in growth media. The B_12_ concentrations in Walne growth medium decreased significantly after autoclaving, resulting in decreased B_12_ concentration in the algal cultures. Although not all hatcheries autoclave vitamins solutions, this practice is relatively common and is worthy of note, as the sharp drop in the B_12_ concentration after autoclaving was significant. The amounts stipulated in media formulations were designed to be in excess for this reason; however, it was not clear if this could nonetheless, reduce the growth rate of *C. calcitrans*, or have subsequent effects upon larval rearing. Previous research has shown that increased B_12_ in f/2 growth medium does not result in increased algal growth rate, as long as minimum B_12_ concentrations are being met (62).

When provided with B_12-_rich diets, Pacific oyster larvae do not deplete or significantly bioaccumulate B_12_ throughout larval development. They appear to maintain similar levels of B_12_ compared to the early D-shelled stage prior to first feeding, thus suggesting that larvae already start out with high B_12_ concentrations derived from non-algal sources. Our hypothesis that differences in B_12_ concentrations of algal feed might be reflected in B_12_ levels in larvae, which would have supported a theory of uptake from dietary sources, was not confirmed in results we report here. Indeed, the B_12_ provided appears to be adequate in all treatments, and while a weak trend toward higher B_12_ concentrations in 10xB_12_ f/2 is seen, the vitamin concentration did not significantly differ from either 15 dpf larvae (closest in size to f/2 larvae) or 19 dpf larvae (closest in age) fed with Walne-grown algae which contained the highest B_12_ concentrations of all treatments, including the B_12_ enriched f/2 medium. These results suggest that larvae are not obtaining their B_12_ primarily from their microalgal feed. With the limited information on B_12_ requirements in bivalve larvae available, however, any potential beneficial effect of B_12_ uptake through diets might be inconsequential as the B_12_ requirements of larvae might have been met by even the lowest B_12_ diets tested. A further decrease in the B_12_ concentration in algal feed could shed additional light on this question; however, a reduction of B_12_ concentration in the growth media could also result in poorly performing algae with unpredictable effects upon larval performance (63).

Higher B_12_ concentrations in the diet did not significantly increase larval growth rates, as seen for the two f/2 algal diets, suggesting that enriching algal feeds with B_12_ alone does not provide an advantage to the aquaculture industry in relation to improving larval growth. Presuming that a minimum level of B_12_ is available, additional B_12_ did not appear to provide any visible benefit, although we did not perform stress or immune challenges to determine if there may, in fact, be benefits for larval survival under adverse conditions. A significant increase in larval growth rate was seen for larvae fed with Walne-grown algae; however, this acceleration of larvae development may be a result of stable temperatures in the 200-L tank compared to the fluctuating lower temperatures in the smaller 20-L tanks (temperature was probably a more important predictor of development rates in bivalves than our treatments (64)). Walne and f/2 seawater enrichments not only vary in B_12_ concentrations, but also in other essential nutrients such as nitrogen. Whether or not the Walne medium improved the quality of the algal diet and eventually benefited larval development was not tested in this study.

A significant decrease in B_12_ concentration was recorded in spat after metamorphosis suggesting that stored B_12_ in larvae was utilized during metamorphosis, or that heavier calcification of the shell increased individual mass relative to soft-tissue mass. The vitamin’s metabolic function during this key life event is unknown. Given the known anti-oxidant properties of B_12_, as well its important role in neurogenesis in other animals, the depletion or dilution of B_12_ reserves during metamorphosis is not surprising. Indeed, metamorphosis appears to be a key life stage wherein B_12_ is important, thus suggesting that further work on larval B_12_ reserves in relation to settlement could be a valuable investigation in relation to hatchery production. For instance, feeding spat for a short duration with a high B_12_ diet after metamorphosis did not replenish the B_12_ concentrations to levels observed before metamorphosis. A more in-depth assessment of larvae and spat prior, during, and post-metamorphosis with different B_12_-containing algal diets might provide further insight into B_12_ sources for larvae and spat. We cannot, however, exclude that the decrease in the vitamin concentration in spat is a consequence of the higher shell to soft-tissue mass ratio in spat compared to larvae. Given that the B_12_ analysis methods are accurate primarily on soft tissue, we cannot exclude this interpretation.

The source of B_12_, whether algal diet or microbial, remains elusive. Our results provide some evidence that B_12_ is sourced from enrichments added to grow the microalgal diet, several other possible sources of B_12_ are worthy of consideration in relation to our findings. For pre-feeding stages, nutrients including essential vitamins might be supplied by maternal egg reserves as previously observed in bivalves and other animals (65-68). Our data, however, indicate that B_12_ concentrations in unfertilised eggs are significantly lower than those of unfed D-shelled larvae, suggesting that another source of B_12_ than their egg reserves is also available to larvae. The trochophore life stage, a free-swimming larval stage prior to shelled D-shaped bivalve larvae, has basic structures such as a mouth, digestive mass (anlagen of the stomach) and intestine, ingesting particles in surrounding water for filter feeing – thus potentially ingesting symbiotic bacteria that could provide B_12_ to the host. Gut microbiome analysis of adult bivalves and whole *Crassostrea* spp. veliger larvae have revealed that the majority of microorganisms are *Protobacteria (*up to ∼95% in larvae) and *Cyanobacteria* ((22) and the references herein (69, 70)). *Cyanobacteria* are known to produce pseudo-vitamin B_12_, the non-bioavailable form of B_12_ for most eukaryotes (59, 71, 72), and are therefore not likely to contribute to bivalve B_12_ uptake. Approximately 45% of the *Proteobacteria*, however are predicted to be B_12_ producers (73), in particular *α-Proteobacteria* and *γ-Proteobacteria* are suggested as B_12_ producers in marine environments (74-77), which are also abundantly present in the microbiota of late *C. gigas* larvae (up to ∼65% in 16 dpf larvae) (70). Thus, various prospective B_12_-producing bacteria are potentially being consumed and digested or colonizing the gut of bivalves providing a stable source of B_12_. This is partially supported by a study in four gastropod species, which found that, of the 270 bacteria strains isolated from the gastrointestinal tract, 87% were B_12_-producing bacteria (78). These authors, however, concluded that only 6% of these bacteria showed high productivity compared to bacteria in surrounding seawater, and none of them were identified as dominant species. Nevertheless, recent studies have shown that microbiomes of oysters can vary throughout larval life (79), under stress conditions (high pH) (80), between hatcheries (81), and rearing locations (82), but larvae and adults usually contain a small core microbiome ((22) and the references herein, (82, 83), which potentially holds the answer to B_12_-producing bacteria symbionts. To shed light on this, further research on oyster larvae is needed, including research into the microbiota of trochophore larvae with a special focus on B_12_-producing species abundance. In addition, the microbiomes of fertilised eggs could also provide some insight into potential vertical or/and horizontal transfer of bacteria from egg to embryo as reported in fish species (84). Furthermore, detection and localisation of cobalamin binding intrinsic factor, a glycoprotein required for uptake of B_12_ that is normally present in the gut of animals - and is predicted in *C. gigas* (GenBank: LOC105331555) - will shed further light on where within the organism the majority of B_12_ uptake occurs for both larvae and adults.

In regards to juveniles (post metamorphosis) and adult bivalves, other potential microbiomes should also be considered as sources of B_12_. Gills are prone to host a variety of microorganisms, wherein previous research has shown that oyster gills can have higher bacterial diversity than the digestive gland, including a large variety of *α-* and *γ-Proteobacteria* (85, 86). Symbiotic relationships between gill bacteria and their bivalve hosts have been well studied in the context, for instance, of chemosynthetic symbioses (87, 88), or the supply of digestion enzymes for celluloses and lignin (89, 90). Vitamin B_12_ concentration in *C. gigas* gills were relatively high, as well as very high in the remaining parts including the gills of the Goolwa cockle, and could therefore indicate potential symbiosis in the gills with B_12_-providing bacteria, or a role of B_12_ in the gills and mantle that deserves further exploration.

## Conclusion

Our data confirms previous results that the Pacific oyster and the commercial scallop tested contain high concentrations of B_12_ with additional evidence that this is also the case for the Goolwa cockle, a species not previously assessed for its B_12_ content. Thus, bivalves provide a reliable source of B_12_ for humans, and can be particularly important in areas of the world where other animal proteins such as meat and dairy products are less available, or people want to reduce the consumption of domesticated farm animals as meat source because of ethical and environmental concerns. Our research, one of the first studies that assessed B_12_ concentrations in different tissues across species, suggests that B_12_ concentrations vary between both species and between tissues within species. Therefore, B_12_ availability in shellfish foods might depend upon which species and which tissue is commonly consumed, for instance scallop adductor muscle - the part of the scallop that is most commonly eaten in both Asian and Western countries - will provide less B_12_ than consuming a whole raw oyster.

The sources of such high levels of B_12_ in bivalves are unknown, as are the metabolic processes that utilise or require them. Differences in B_12_ tissue accumulation suggest differences in physiological functions for B_12_ among different bivalve species. Further analysis of different tissue types in additional bivalve species will provide better insight into the possibility that differences in B_12_ accumulation in tissues are conserved across different families or habitats, and whether or not there are patterns in freshwater *versus* marine species, or based upon spatial habitat distribution, such as levels between bivalves in intertidal or sublittoral zones. Such information could provide better insight in the origin of B_12_ in bivalve species, and the reason for such high levels of accumulation. Given that oysters are filter feeders ingesting a large variety of microorganisms and microalgae, we hypothesised that B_12_, which can only be produced *de novo* by bacteria, could be produced by the animal’s own gut or gill microbiome, such is the case in terrestrial ruminants. Alternatively, it was possible that B_12_ is ingested by filter feeding on bacteria contained in particulate matter, or by ingesting B_12_-rich microalgae, with B_12_ assimilated into the algae through known bacteria-algae symbiosis. Our results from the feeding experiments, however, did not confirm an increase in B_12_ concentration in oyster larvae when fed with B_12_-enriched algal feed, thus suggesting that algal diet might not be the main source of B_12_ (at least when high amount of B_12_ is provided). In addition, unfed D-shelled larvae already contained high amount of B_12_, thereby implying B_12_ uptake was provided by symbiotic microorganisms potentially acquired during the early trochophore larval stage. However, B_12_ concentrations in the digestive tract of adult oysters are one of the lowest of all oyster tissues, indicating that B_12_ production of a potential gut microbiome might be low. Overall, we could not unambiguously reach a clear conclusion about the source of B_12_ in oysters and our experience indicates that identifying the B_12_ origin in oysters is not a simple task. Further investigations of core microbiome communities in the gut and other tissues such as gills in relation to known B_12_-producing bacteria, as well as experiments involving elimination of B_12_ through use of B_12_ inhibitors might shed more light on the question of the origin of the high levels of B_12_ in bivalves. Our results provide valuable information in relation to basic oyster larvae physiology as well as aquaculture applications. Vitamin B_12_ concentrations are stable throughout oyster larval development, but data suggested a high requirement of B_12_ during metamorphosis. Understanding how B vitamins are assimilated and used metabolically is interesting, particularly in light of genetic disorders in humans that lead to deficiencies (e.g. pernicious anaemia). It is also important for livestock and finfish diets, where vitamin supplementation is common to optimize growth rates. Although shellfish diets in hatcheries are not currently supplemented with vitamins beyond what is provided to the algae in growth media, investigations such as this study are important to assess vitamin requirements of larvae during hatchery production. We can conclude that although B_12_ concentrations in microalgal cultures can be increased by supplementing growth media with additional B_12_, this does not necessarily lead to an increase in B_12_ concentrations within larvae nor does it lead to faster growth. Whether B_12_ supplementation however, might provide some health benefits to larvae based upon increased survival when challenged with pathogens, or whether it increases rates of metamorphosis still needs to be determined. Based on our results, we recommend caution when autoclaving as sterilisation of growth media after vitamins have been added as this clearly leads to a decrease in B_12_ concentrations in microalgal cultures.

## Acknowledgments

For help with rearing of Pacific oyster larvae, we would like to thank Mark Gluis, Dr Penny Ezzy-Miller and the hatchery team from SARDI.

## Supplementary Information

**S1 File: Sample information, weight and B**_**12**_ **concentrations**

**S2 File: Media composition**

